# Molecular mechanism of quorum sensing inhibition in *Streptococcus* by the phage protein paratox

**DOI:** 10.1101/2021.06.03.446943

**Authors:** Nicole R. Rutbeek, Hanieh Rezasoltani, Trushar R. Patel, Mazdak Khajehpour, Gerd Prehna

## Abstract

*Streptococcus pyogenes*, or Group A Streptococcus, is a Gram-positive bacterium that can be both a human commensal and pathogen. Central to this dichotomy are temperate bacteriophages that incorporate into the bacterial genome as a prophage. These genetic elements encode both the phage proteins as well as toxins harmful to the human host. One such conserved phage protein paratox (Prx) is always found encoded adjacent to the toxin genes and this linkage is preserved during transduction. Within *Streptococcus pyogenes,* Prx functions to inhibit the quorum-sensing ComRS receptor-signal pair that is the master regulator of natural competence, or the ability to uptake endogenous DNA. Specifically, Prx directly binds and inhibits the receptor ComR by unknown mechanism. To understand how Prx inhibits ComR at the molecular level we pursued an X-ray crystal structure of Prx bound to ComR. The structural data supported by solution X-ray scattering data demonstrate that Prx induces a conformational change in ComR to directly access the DNA binding domain. Furthermore, electromobility shift assays and competition binding assays reveal that Prx effectively uncouples the inter-domain conformational change that is required for activation of ComR by the signaling molecule XIP. Although to our knowledge the molecular mechanism of quorum-sensing inhibition by Prx is unique, it is analogous to the mechanism employed by the phage protein Aqs1 in *Pseudomonas aeruginosa.* Together, this demonstrates an example of convergent evolution between Gram-positive and Gram-negative phages to inhibit quorum-sensing, and highlights the versatility of small phage proteins.

## INTRODUCTION

*Streptococcus pyogenes* is responsible for a broad range of human diseases, ranging from minor skin infections and pharyngitis, to more serious complications, such as rheumatic fever and toxic shock syndrome (1, 2). A significant contributor to these varying levels of pathogenicity is the presence, or absence, of bacteriophage encoded toxin and virulence genes (3). These genes are actively spread between different strains and different species of *Streptococcus* through horizontal gene transfer. This process commonly occurs by direct infection of phage, but also includes the mechanisms of transduction and natural competence (transformation) (4, 5).

The temperate bacteriophage and phage like elements associated with *S. pyogenes*, are dsDNA phage belonging to the *Siphoviridae* family (6, 7). This includes the lysogenic T12 and SF370 phages that can exist as a prophage within a bacterial host. These prophages consist of stable genetic elements inserted into a Streptococcal genome (6). There are often multiple prophages present within the genomes of each strain of *S. pyogenes*, many of which encode the deadly toxin and virulence genes characteristic of Group A Streptococcus (GAS) infections. These include the superantigen SpeA, a key toxin that results in scarlet fever and Streptococcal toxic shock syndrome, and enzymes such as phospholipases and DNases, which also contribute significantly to *S. pyogenes* virulence (8, 9). However, given the right conditions the prophage can exit through the lysogenic cycle and self-excise from the bacterial genome to produce mature phage that can infect a new host (6). After infection, the mature bacteriophage can then incorporate as a prophage and potentially create a new virulent strain of *S. pyogenes.* In line with this ability, bacteriophages are thought to be the primary force driving clonal diversity in GAS (7, 10).

In addition to phage infection, many species of *Streptococcus* acquire exogenous DNA through natural competence. For Gram-positive bacteria, natural competence (natural transformation) is a quorum sensing regulated process in which bacteria control the expression of a number of genes that encode the machinery for both the acquisition and the incorporation of DNA (4, 11, 12). Although natural transformation is difficult to observe in *S. pyogenes* (13, 14), it contains all the genes necessary for natural competence, including the ComRS quorum sensing pathway (14). ComR is a quorum sensing receptor and a transcription factor that belongs to the Rgg sub-group of the RRNPP protein family, many of which regulate virulence in pathogenic *Streptococcus* species (12). ComS is a secreted small peptide pheromone that when processed into its mature form of ~7-8 amino acids termed XIP (<u>s</u>igX inducing <u>p</u>eptide), it is able to bind and activate ComR (15). The activated ComR:XIP complex recognizes both the *comS* (XIP) and the alternative sigma-factor *sigX* promoter regions (15), creating a positive feedback loop for the quorum sensing circuit. SigX itself binds to CIN-box promoters which leads to the expression of late genes that are required for natural competence (16, 17).

Previous work to understand the role of ComRS quorum sensing in *S. pyogenes* made the discovery that expression of the small prophage protein paratox (Prx) is also induced in a ComRS dependent manner (15, 18). Specifically, Prx can be directly induced by XIP since its expression is under the control of a CIN-box promoter (18). Prx is highly conserved and found in the genomes of *S. pyogenes* as well as many other species of *Streptococcus,* including *S. agalactiae* and *S. suis* (18). Interestingly, *prx* is always encoded at the 3’ end of the prophage and is found adjacent to a toxin or virulence gene, such as *speA* (18, 19). Furthermore, *prx* and the adjacent toxin are in ‘linkage disequilibrium’ such that they remain together as one genetic cassette during homologous recombination and when the bacteriophage exits the lysogenic cycle (19).

While the biological role of Prx relative to its genetically paired toxin remains to be determined, Prx has been shown to function as a negative regulator of natural competence in *Streptococcus.* Specifically, Prx binds directly to the apo-form of ComR preventing DNA binding *in vitro* and acts as a repressor of the *S. pyogenes* competence regulon *in vivo* (18). Together this demonstrates not only a link between transduction and natural competence, but also may help to explain in part why *S. pyogenes* is observed to have low levels of natural transformation.

Although Prx inhibits ComRS dependent quorum sensing in *Streptococcus*, the biochemical and molecular mechanism of this interaction is unknown. It was recently found that the small phage protein Aqs1 inhibits the general quorum sensing pathway in the Gram-negative bacteria *Pseudomonas aeruginosa* by binding the transcription factor LasR (20). However, Prx and ComR are not structurally related to either Aqs1 or LasR respectively. Prx has a unique fold with distant similarities to the lambda phage head-tail joining protein GpW (18, 21) and ComR undergoes a drastic conformational change upon binding XIP (22, 23). ComR is a monomer that contains a DNA binding domain (DBD) packed against a tetratricopeptide repeat (TPR) domain. The monomer conformation is such that several conserved residues in the DBD are shielded and partially buried to prevent interaction with DNA (22, 23). Once XIP binds the TPR domain this results in an extensive conformational change that both frees the DBD and allows ComR to dimerize using both the TPR domain and a DBD domain-swap for interaction with DNA (22). As our past work has shown Prx can interact with apo-ComR (18), Prx could be exerting its inhibitory effect to block any or multiple steps in the XIP dependent activation of ComR.

To elucidate the molecular mechanism of inhibition of the ComRS quorum sensing pathway by Prx, we determined high-resolution structures of a ComR:Prx complex. Our crystallographic data supported by small-angle X-ray scattering (SAXS), demonstrates that Prx induces a conformational change in apo-ComR to release the DNA binding domain from the TPR domain so that Prx can directly bind the DNA-interacting residues. Additionally, electromobility shift assays (EMSA) combined with fluorescently labeled peptide binding experiments show that Prx interacts with ComR independently of XIP and that Prx can form a ternary complex with ComR:XIP. Our results not only reveal a dynamic mechanism of quorum-sensing and transcription factor inhibition, but also demonstrate an example of convergent evolution between Gram-positive and Gram-negative phages. Specifically, the strategy of phages to modulate the pathways of their hosts as a means of self-preservation using highly versatile small bacteriophage proteins.

## RESULTS

### Prx binds each known class-type of ComR protein

Although Prx is highly conserved (18), ComR shows high residue variability across Streptococcal strains and species. Namely, the amino acid conservation of the ComR TPR domain can diverge significantly, especially within the XIP binding pocket (23). This is attributed to the fact that there are three classes of ComRS pathways. Type I ComR proteins (Salivarius group) recognize hydrophobic XIP pheromones, Type II ComR proteins (Bovis, Mutans, Pyogenic groups) recognize XIP pheromones with a WW-motif, and Type III ComR proteins (Suis group) recognize XIP pheromones with a W-XX-W-motif (14, 24, 25). Moreover, each individual ComR exhibits a unique profile of XIP specificity ranging from only recognizing their specific species XIP (*S. mutans)* to being able to be activated by almost any XIP (*S. bovis*) (23).

Given the variability of ComR proteins, we asked if Prx has a clear specificity for any of the ComR orthologs. Using purified *S. pyogenes* MGAS315 Prx we performed binding assays with Type II ComR proteins (*S. pyogenes* MGAS5005 and *S. mutans*), as well as a Type I ComR (*S. thermophilus*) and a Type III ComR protein (*S. suis*). It is important to note that ComR MGAS315 could not be successfully purified. Prx interacts with each type of ComR in a pull-down assay (Fig. 1A) and forms a stable complex with each ComR ortholog as demonstrated by size-exclusion chromatography (SEC) (Fig. S1). While qualitative, the pull-down assay appears to suggest that Prx may have a preference for Type II ComR proteins. However, given that ComR:Prx complexes can be isolated by SEC for each ComR ortholog, we conclude that Prx proteins can recognize all three known class-types of ComR proteins, indicating a conserved mode of interaction. This broad recognition is underscored by the fact that *S. mutans* lacks any known *prx* genes (18).

**Figure 1.**
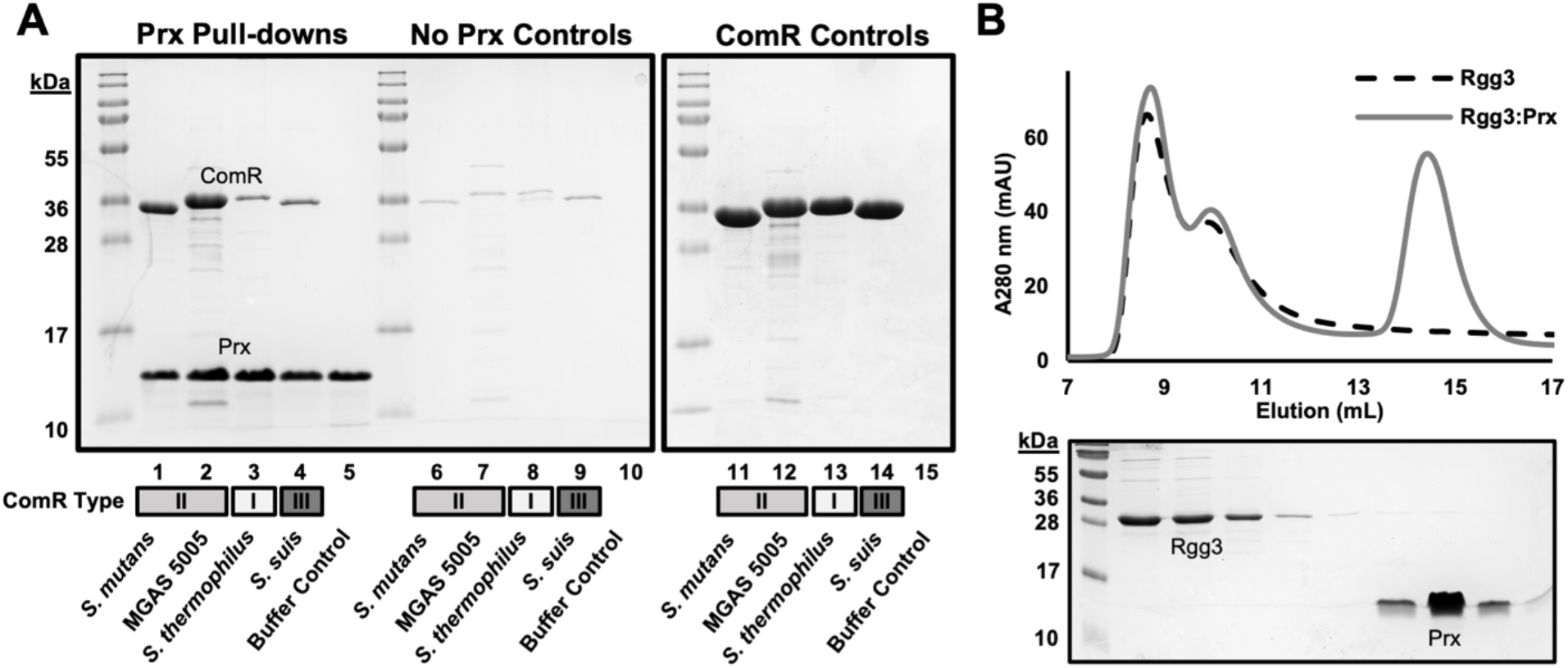
Prx forms stable complexes with various ComR orthologs. A) Pull-down assay of Prx with various ComR orthologs including type II ComR proteins (ComR *S. mutans,* ComR MGAS5005) type I (ComR *S. thermophilus*) and type III (ComR *S. suis*). Controls with beads without Prx (no Prx Controls) and purified protein (ComR controls) are shown. B) Size exclusion chromatography (SEC) binding assay with the ComR structural relative Rgg3. The top panel shows the chromatogram and the bottom panel a Coomassie stained SDS-PAGE gel of the corresponding fractions.

ComR shares high structural homology to the proteins that are members of the Rgg sub-family of RRNPP peptide signaling systems (23, 26). Like ComR, Rgg-subfamily proteins possess a DBD and TPR domain, form dimers, and have their activity modulated by small peptide pheromones (12, 27). Given that other Rgg proteins are also found in *Streptococcus,* we asked if Prx could bind a close structural relative. To test this, we assessed the ability of ComR to interact with Rgg3 from *S. pyogenes* by SEC. However, there was no noticeable shift in elution volume to indicate complex formation or observed co-elution of Prx with Rgg3 as visualized by SDS-PAGE gel (Fig. 1B). It is possible that a certain set of conditions may be required for the interaction of Prx with Rgg3; however this is unlikely as Prx can readily recognize apo-ComR. With the biological role of Rgg3 in the quorum sensing regulation of virulence and biofilm formation in *Streptococcus* (12), this suggests that Prx has specifically evolved to inhibit quorum sensing regulated natural competence in the bacterial host.

### Prx interacts directly with the ComR DNA binding domain

Our goal was to visualize the molecular interaction between ComR and Prx using X-ray crystallography, however we were not able to crystalize a full-length ComR:Prx complex with the ComR orthologs purified in Figure 1. In order to help delineate the ComR:Prx interaction surface, we assayed the ability of Prx to bind to either the DBD or TPR domain of ComR. Using SEC, we were able to determine that Prx binds directly to the DBD and does not form a stable complex with the TPR domain construct (Fig. 2A, Fig. 2B, and Fig. S2).

**Figure 2.**
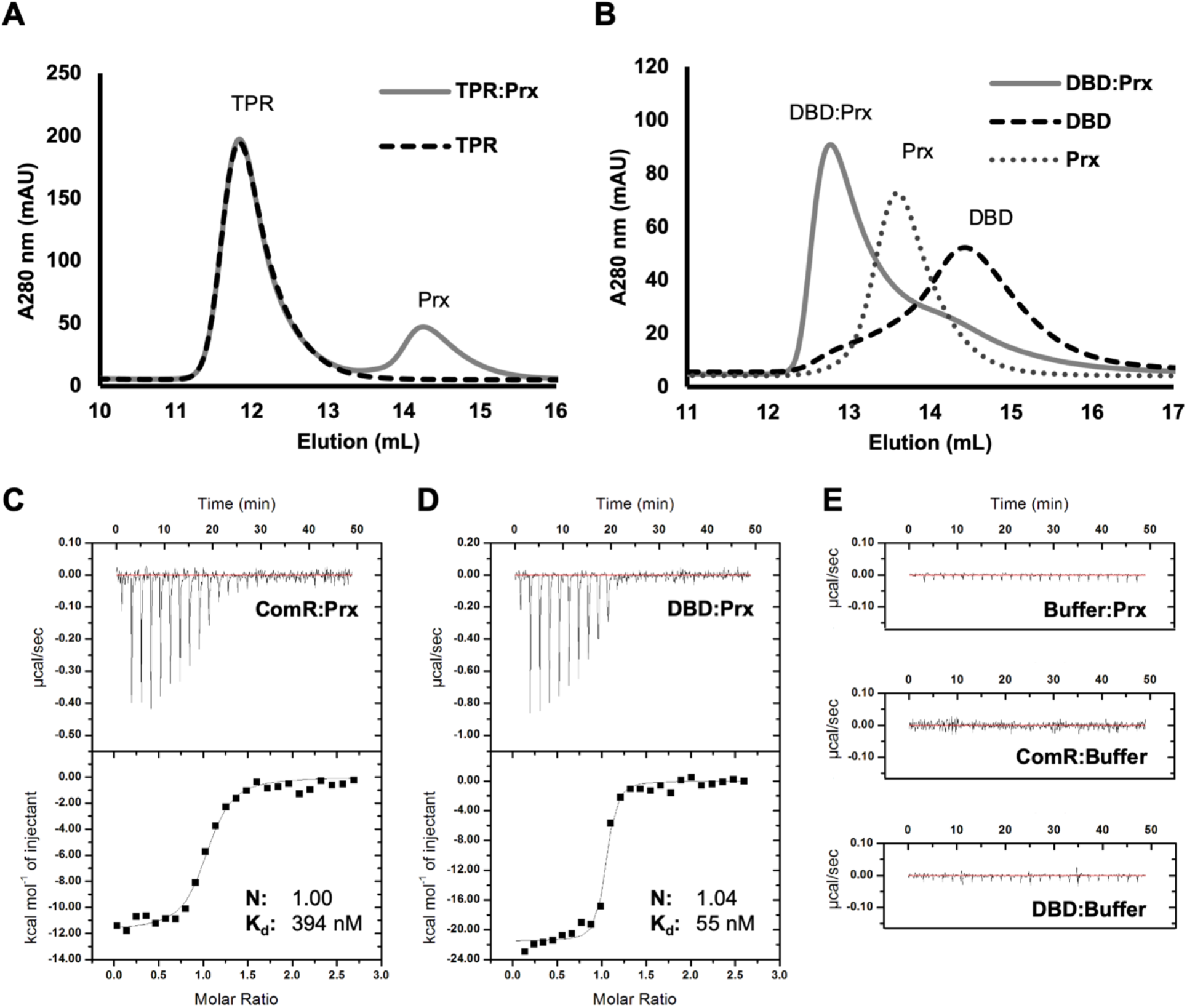
Prx binds the ComR DBD with high affinity. A) Size exclusion chromatography binding assays of Prx with the TPR domain (left panel) and the DBD (right panel) of ComR *S. mutans.* ComR minimal domains TPR or DBD incubated with Prx are shown in grey, the TPR or DBD alone as black dashed lines, and Prx alone as a dotted grey line. B) Isothermal titration calorimetry was performed with Prx and full length ComR and in C) with Prx and the ComR DBD. D) Isothermal titration calorimetry controls of Prx, full-length ComR, and the ComR DBD with buffer. Prx forms a 1:1 stoichiometric complex with both full-length ComR and the ComR DBD with a binding constant of 392 nM and 52 nM respectively.

The interaction of Prx with the DBD was further characterized using isothermal titration calorimetry (ITC). Similar to past results, we observed that Prx binds apo-ComR *S. mutans* with sub-micromolar affinity (Fig. 2C) (18). However, Prx was able to bind the minimal DBD construct with significantly higher affinity than the wild-type ComR protein construct (Kd of 52 ± 15 nM compared to 392 ± 82 nM) (Fig. 2C and Fig. 2D). ITC binding controls are shown in Figure 2E. From these ITC curves, the thermodynamic parameters for the formation of ComR:Prx complex (ΔH= −11.8 ± 0.3 kcal/mol and ΔS = −10 ± 1 cal/mol/K) and the DBD:Prx complex (ΔH= −20.8 ± 0.3 kcal/mol and ΔS = −37 ± 1 cal/mol/K) were determined. The favorable ΔH and unfavorable ΔS is an indicator that the ComR:Prx interaction is likely electrostatic in nature and stabilized by hydrogen-bonding with minimal hydrophobic interactions. Furthermore, the magnitudes of the enthalpy and entropy of formation for the ComR:Prx complex are significantly smaller than that of the DBD:Prx complex. The less favorable enthalpy and more favorable entropy of Prx binding apo-ComR as compared to the DBD, could be indicative of an induced conformational change upon interaction with full-length ComR. As ComR undergoes significant structural changes of the DBD relative to the TPR for activation (23), a potential Prx induced conformational change could be by-passed with just using the minimal DBD construct.

### Prx contacts residues critical for the interaction of ComR with DNA and for the stabilization of the ComR apo-conformation

Despite not being able to obtain Prx in complex with full-length ComR, we were able to co-crystallize Prx with the DBD of ComR. DBD:Prx co-crystals were obtained with both purified minimal DBD bound to Prx and by *in situ* proteolysis with full-length ComR complexed with Prx (Fig. 3, Fig S3, and Table 1). The crystals grew in similar conditions and both diffracted to 1.65 Å and 1.60 Å, respectively. Data collection and refinement statistics are presented in Table 1. The obtained complexes are nearly identical except for differences in space group (P2_1_2_1_2_1_ and P4_1_2_1_2), and their cloning and protease digestion artefacts (Fig. S3A). Additionally, comparing the conformations of both the DBD and Prx observed in the co-crystal complex to the DBD of apo-ComR (PDBid: 5FD4) and Prx alone (PDBid: 6CKA), shows no significant variation (Fig. S3B).

**Table 1.** X-ray data collection and refinement statistics.

|  | DBD:Prx | DBD:Prx (proteolysis) |
| --- | --- | --- |
| **Data Collection** |  |  |
| Wavelength (Å) | 1.28329 | 0.97911 |
| Space group | P2<sub>1</sub>2<sub>1</sub>2<sub>1</sub> | P4<sub>1</sub>2<sub>1</sub>2 |
| Cell dimensions |  |  |
| *a*, *b*, *c* (Å) | 38.56 41.92 90.0 | 36.8 36.8 189.3 |
| $\alpha$, $\beta$, $\gamma$ (°) | 90.0 90.0 90.0 | 90.0 90.0 90.0 |
| Resolution (Å) | 45 -1.65 (1.68-1.65) | 36.14-1.60 (1.66-1.60) |
| Total reflections | 111682 (5567) | 215198 (7232) |
| Unique reflections | 18234 (918) | 18256 (1746) |
| CC(1/2) | 0.998 (0.867) | 0.999 (0.641) |
| *R*<sub>merge</sub> | 0.067 (0.727) | 0.09 (1.30) |
| *R*<sub>pim</sub> | 0.029 (0.316) | 0.03 (0.46) |
| *I*/σ*I* | 15.2 (2.3) | 15.6 (1.5) |
| Completeness (%) | 99.9 (100.0) | 100.0 (99.7) |
| Redundancy | 6.1 (6.1) | 11.7 (8.5) |
| **Refinement** |  |  |
| *R*<sub>work</sub> / *R*<sub>free</sub> (%) | 18.3 / 20.4 | 17.9 / 20.4 |
| Average B-factors (Å<sup>2</sup>) | 31.7 | 25.7 |
| Protein | 30.3 | 24.1 |
| Ligands | - | 53.6 |
| Water | 39.8 | 32.1 |
| No. atoms | 1250 | 1312 |
| Protein | 1062 | 1091 |
| Ligands | 0 | 32 |
| Water | 188 | 206 |
| Rms deviations |  |  |
| Bond lengths (Å) | 0.004 | 0.005 |
| Bond angles (°) | 0.660 | 0.690 |
| Ramachandran plot (%) |  |  |
| Total favored | 100.00 | 99.24 |
| Total allowed | 0.00 | 0.76 |
| PDB code | 7N10 | 7N1N |

Statistics for the highest-resolution shell are shown in parentheses

**Figure 3.**
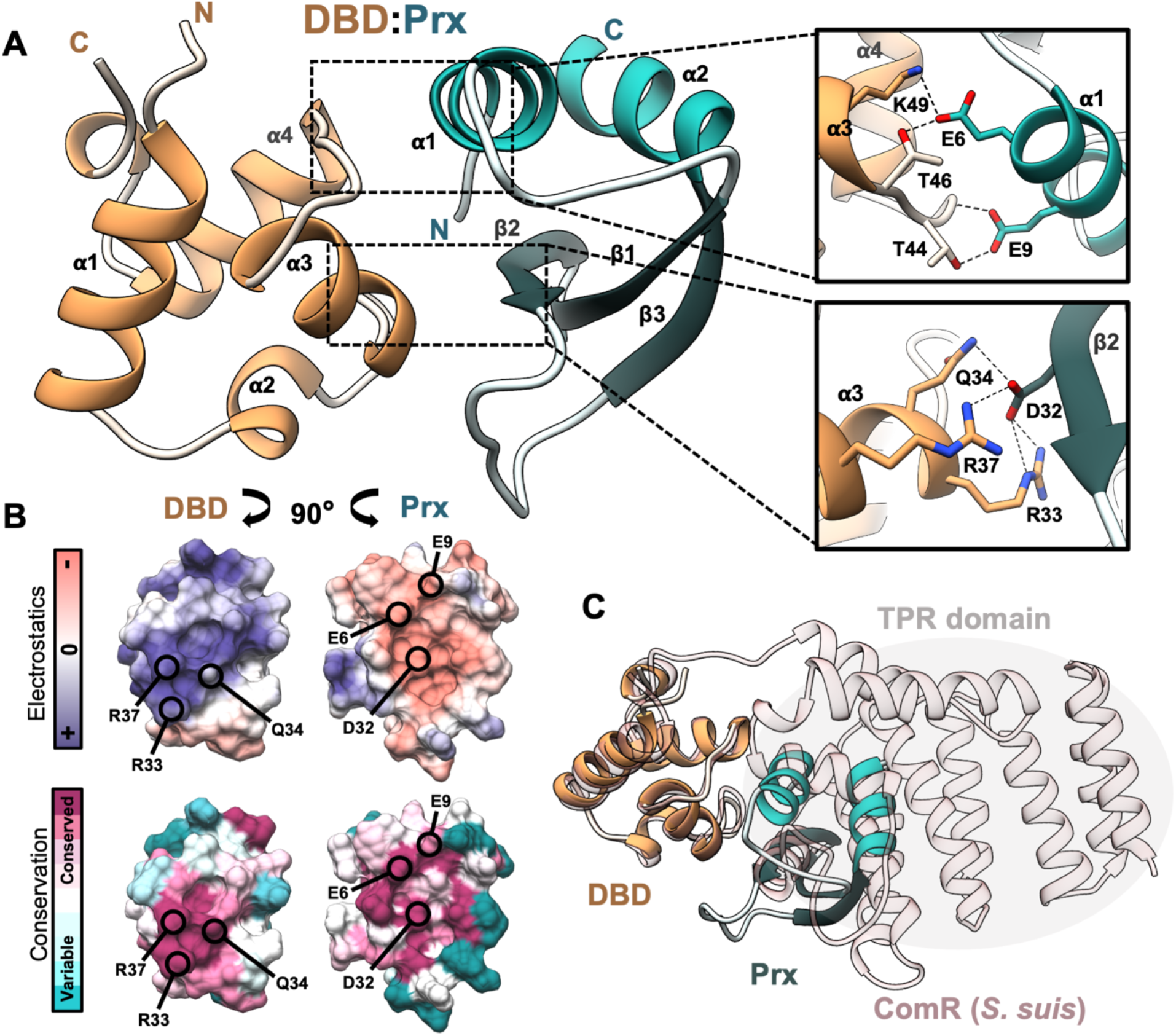
Co-crystal complex of Prx and the DNA binding domain of ComR. A) Co-crystal complex of the minimal ComR DBD (orange) with Prx (cyan). Inset boxes demonstrate key hydrogen-bond and salt-bridge interactions (dashed-lines) between conserved residues in Prx and the residues in the DBD. The chosen residues in the DBD are critical for recognizing DNA and stabilizing the inactive apo-ComR conformation. Secondary structures are labeled by alpha-helix (α) and beta-strand (β) with the N-terminus and C-terminus labeled for each protein. B) Molecular surface representations showing the electrostatics (top) and residue conservation (bottom) of the DBD and of Prx. The represented views are the structures in panel A rotated an opposing 90 degrees to display the surfaces in each protein at the interaction face. Select residues from panel A are indicated. C) Overlay of the DBD:Prx complex on the structure of full-length ComR from *S. suis* (light pink) (PDBid: 5FD4) aligned by the DBD showing a steric clash with the TPR.

Analysis of the DBD:Prx complex interface reveals that Prx directly interacts with the DNA binding surface of ComR (Fig. 3A). The interaction is highly electrostatic in nature, with Prx using a large negative surface consisting of conserved acidic residues to complement the conserved positive DNA-interaction face of the ComR DBD (Fig. 3B and Fig. S3C). At this interface, Prx makes several hydrogen bond and salt-bridge contacts with key residues in the DBD that are critical for interaction with DNA. Specifically, Prx utilizes D32 to make hydrogen bonds with both the essential arginine residues R33 and R37 in ComR that are responsible for DNA recognition (22) (Fig. 3A). Furthermore, residue D32 is the focal point for a stabilized hydrogen bonding network that positions Prx E44 to make a salt-bridge with ComR R33, Prx D38 to salt-bridge with ComR R37 and for the arginine sidechains to be further stabilized by hydrogen bond interactions with the Prx mainchain. This binding network is extended across the full face of the DBD:Prx interaction by Prx E6 making a hydrogen bond with the invariable ComR residue K49 and Prx E9 providing a mainchain hydrogen bond with ComR T44 (Fig. 3A).

Interestingly, Prx D32 also makes direct contact with ComR Q34 as part of the extensive hydrogen bonding network. This is especially important as ComR Q34 (Q40 in *S. suis*) makes hydrogen bond contacts with conserved residues in the ComR TPR. This interdomain-contact is required to hold the DBD against the TPR to stabilize the apo-ComR conformation in its inactive form (23). If the DBD:Prx complex is aligned to apo-ComR using the DBD, we observed that Prx is unable to interact with the DNA binding residues of ComR due to a large steric clash as this surface is held tightly against the TPR domain (Fig. 3C). As such, to bind ComR, Prx must induce a conformational change in apo-ComR to interact with the DBD. Based on this, Prx likely contacts Q34 as part of the mechanism to release the DBD from the TPR. Overall, this model is supported by the ITC binding data which shows a loss in ΔH and a gain in ΔS towards the binding energy relative to that of the minimal DBD construct (Fig. 2). Namely, a cost in ΔH to break the ComR DBD:TPR interaction surface and a gain in rotational freedom in ΔS as the DBD is released.

To verify the observed important structural contacts in the DBD:Prx complex and to support our structural model, we performed site-directed mutagenesis to probe the interaction surface (Fig. 4 and Fig. S4). Multiple Prx protein variants were assessed for their ability to bind to full length ComR using SEC. Prx variants of the three residues highlighted in Figure 3A (PrxE6A, PrxE9A, PrxD32A) were unable to form a complex with ComR, while Prx D12A which is not at the DBD interface was still able to form a complex (Fig. 4A, Fig. 4B, and Fig. S4). Additionally, Prx residue F31 which forms van der Waals at the DBD:Prx interface was also substituted to alanine but had no effect on binding. As an additional control, we collected circular dichroism (CD) spectra to verify that the observed disruptions in complex formation where not due to misfolding of the Prx variants. As shown in Figure 3C, the collected CD spectrum of each variant closely matched the wild-type Prx indicating that the loss of function in each variant is due to specific molecular contacts. Furthermore, given that SEC selects for high affinity interactions, we also tested the ability of Prx D32A to bind ComR using ITC to see if the K_d_ is simply reduced. As shown in Figure 4D, the interaction with ComR is completely disrupted with the substitution of Prx D32A.

**Figure 4.**
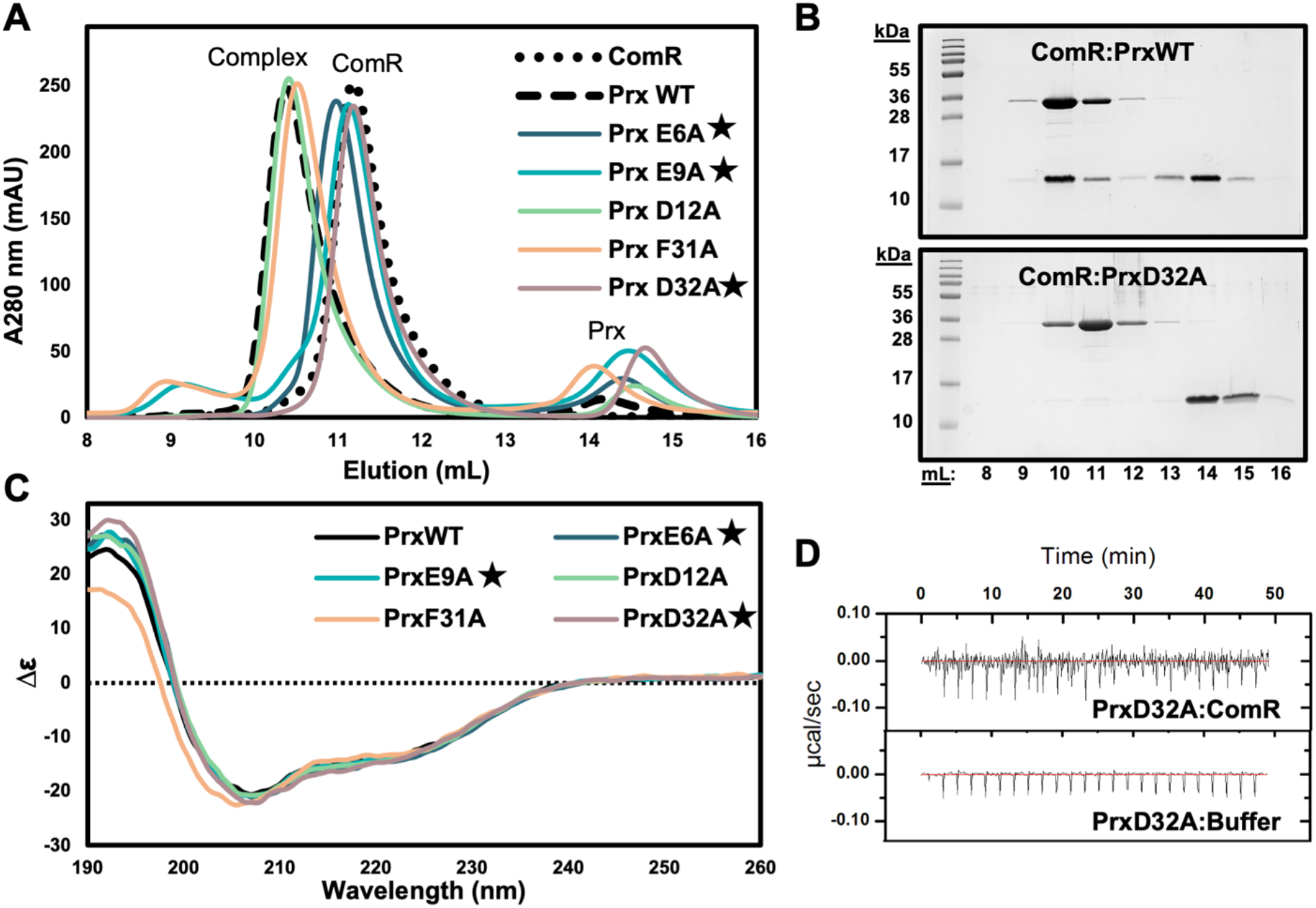
Conserved acidic residues in Prx are required for interaction with ComR. A) Size exclusion chromatography binding assays of Prx variants with wild-type ComR. The chromatogram of ComR alone is indicated by a dotted black line and the chromatogram of ComR incubated with wild-type Prx as a dashed black line. Each Prx variant assay is plotted in color. Starred residues indicate contacts shown in Figure 3A that when mutated result in a loss of complex formation. B) Representative Coomassie stained SDS-PAGE gels of SEC binding controls and a Prx protein variant with ComR. C) Circular Dichroism spectra controls of the Prx variants compared to wild-type Prx. The Prx variant color key and labeling are matched with panel A. D) ITC experiment and control for PrxD32A and full length ComR.

### Small-angle X-ray scattering (SAXS) data provides a model for the full-length ComR:Prx complex

Since our complex structure includes only the DNA binding domain of ComR, we used a SEC-coupled SAXS set-up to visualize the full ComR:Prx complex. SAXS provides low-resolution structural information that is useful for visualizing the overall shape of protein complexes in solution, including large conformational changes (28). SEC-SAXS datasets were collected for ComR, Prx and the ComR:Prx complex, each in the same buffer (gel filtration buffer) at a concentration of 9 mg/mL (Fig. S5A). Sample selection and buffer subtraction was performed using CHROMIXS (29) and the datasets were processed and analyzed with PRIMUS (30) and ScÅtter (www.bioisis.net).

A plot of scattering intensity vs. scattering angle from the buffer-subtracted SEC-SAXS data and the subsequent Guinier fit for each sample is shown in Figure 5A. The linear fit of the data in the low-q region as shown in the each Guinier plot indicates mono-disperse protein samples. The fitted data in the Guinier regions provided Rg (radius of gyration) and I(0) values (Table 2), corresponding to molecular weights that are near the theoretical molecular weights of 35 kDa and 43 kDa for ComR and ComR:Prx respectively. In agreement with past SEC analysis, the hydrodynamic radius of Prx appears larger than expected corresponding to a molecular weight near 16 kDa (18). We further processed the SAXS data, performing Kratky analysis (Fig. 5B) and Porod analysis (Fig. S5B) to assess the globular nature of each protein. As shown in the dimensionless Kratky plot (Fig. 5C), the proteins are overall globular, while both Prx and the ComR:Prx complex appear to be slightly more flexible in solution. This is further demonstrated by the Porod exponent (P_X_), determined by fitting a linear curve to the Porod region of the Porod-Debeye plot, where a completely globular protein would have a P_X_ of 4 and a completely flexible or disordered protein would have a P_X_ of 2 (31, 32). ComR is globular, with a P_X_ of approximately 3.9, and ComR:Prx and Prx, while still globular are slightly more flexible with Porod exponents of 3.7 and 3.6 respectively.

**Table 2.** Small-angle X-ray scattering data.

|  | ComR | ComR:Prx | Prx |
| --- | --- | --- | --- |
| **Guinier Analysis<sup>a</sup>** |  |  |  |
| Rg (Å) | 23.59 ± 0.03 | 27.83 ± 0.04 | 16.10 ± 0.05 |
| I (0) | 0.074 ± 5.1 x10<sup>-5</sup> | 0.085 ± 7.0x10<sup>-5</sup> | 0.020 ± 3.0x10<sup>-5</sup> |
| **Porod Analysis<sup>b</sup>** |  |  |  |
| Porod Exponent (P<sub>x</sub>) | 3.9 | 3.7 | 3.6 |
| Porod Volume (Å<sup>3</sup>) | 61460 | 76200 | 17450 |
| **P(r) Distribution<sup>a</sup>** |  |  |  |
| D<sub>max</sub> (Å) | 78.0 | 98.8 | 43 |
| Rg (Å) | 23.5 ± 0.07 | 28.3 ± 0.09 | 15.4 ± 0.02 |
| P(r) quality of fit | 0.823 | 0.772 | 0.891 |

<sup>a</sup>Data analysis in PRIMUS <sup>b</sup>Data analysis in ScÅtter IV

**Figure 5.**
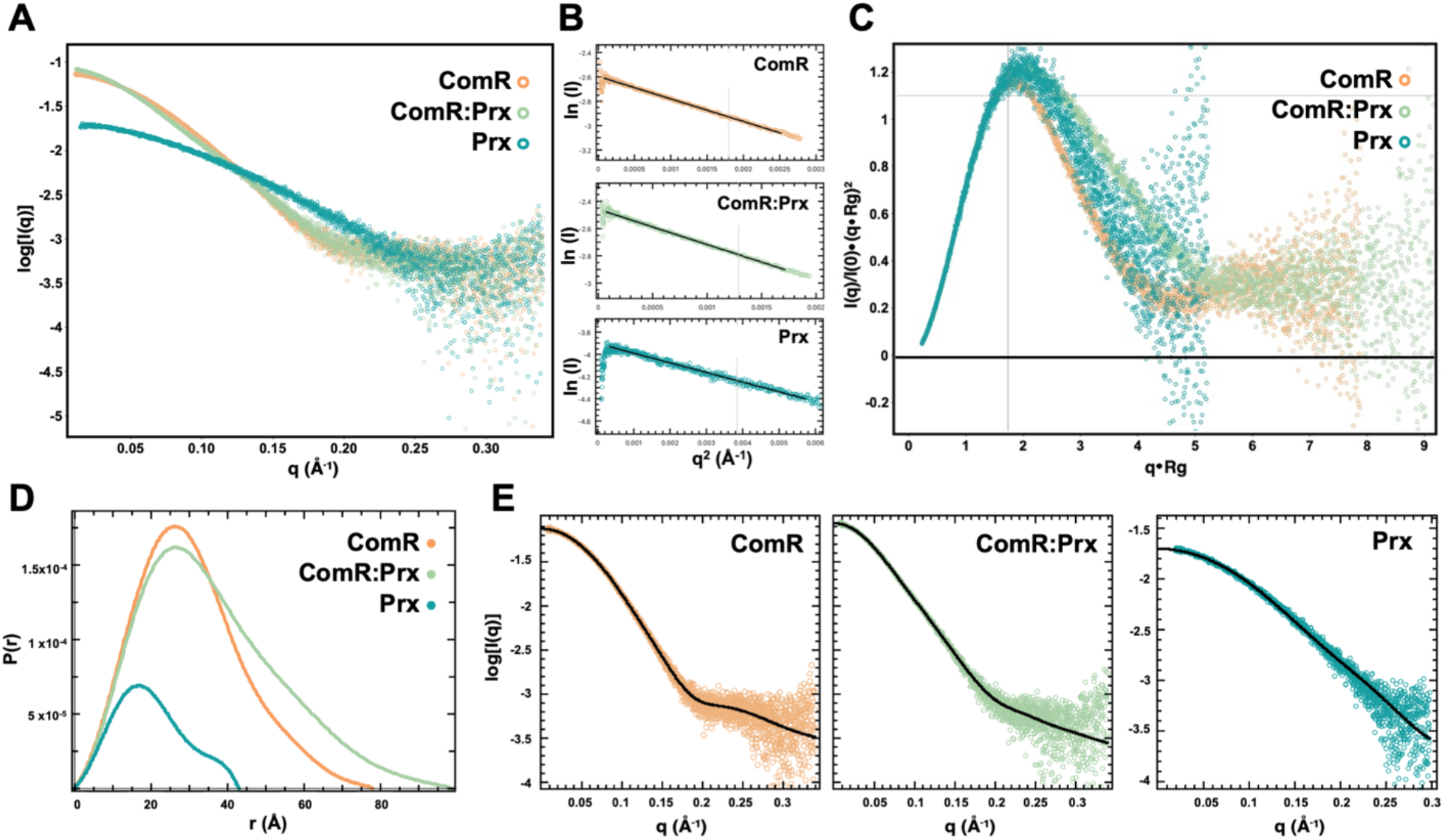
Small-angle X-ray scattering characterization of Prx and ComR. A) Scattering intensity (log I(q)) plotted against scattering angle (q(Å)). B) Guinier fit of each experiment in panel A. C) Normalized Kratky plots for ComR, ComR:Prx, and Prx demonstrating relative flexibility. D) Distance distribution plot (P(r)) for the determination of Rg and D_max_ for ComR and ComR:Prx. E) Fit of the calculated P(r) distribution to the scattering data. Each plot is colored by protein sample, ComR (orange), ComR:Prx (light green), Prx (blue).

As the ComR, ComR:Prx, and Prx samples are homogenous, we determined their real-space electron pair distribution function, or P(r), using GNOM (33) (Fig. 5D and Fig. 5E). The P(r) distribution yielded Rg values of 23.5 ± 0.07 Å for ComR, 28.5 ± 0.09 Å for ComR:Prx, and 15.5 ± 0.02 Å for Prx which agree with the Rg values derived from Guinier analysis (Table 2). Following the calculation of the P(r) distribution, the corresponding scattering data was then used for the estimation of low-resolution electron density maps by DENSS (34). We also utilized *ab initio* modeling program DAMMIN (35),and DAMAVER to obtain low-resolution envelope. The chi (*X*) values of ~1.12 (ComR), ~1.1 (ComR:Prx) and ~1.04 (Prx) suggest an excellent agreement between SAXS data and DAMMIN models derived data. Furthermore, the normalised special discrepancy (NSD) values of 0.49 (ComR), 0.56 (ComR:Prx) and 0.61 (Prx) demonstrate that individuals models are highly similar with each other for each sample. We aligned the structures of apo-ComR, a proposed ComR:Prx, and Prx into their respective SAXS density maps using Chimera (36). The resulting models for ComR and ComR:Prx are shown in Figure 6. As presented in Figure 6A and 6B, DAMMIN models matched closely to the maps provided by DENSS. Additional views of the density fits are displayed in Figure S6A and Figure S6B. The models for Prx are shown in Figure S6C and Figure S6D. ComR *S. suis* fits well into the calculated density map of ComR *S. mutans*, with a smaller portion on one end that corresponds to the DBD, and a larger section corresponding to the TPR domain. For the ComR:Prx complex we observe a significant increase in density reflecting Prx binding to the DBD and releasing it from the TPR domain. As such, the TPR domain of ComR (*S. suis*) was placed in the SAXS envelope with the DBD:Prx complex modeled into the remaining density. Given that density for the DBD:Prx was clearly visible, it is likely that even when released the DBD may have preferred orientations relative to the TPR domain when in complex with Prx. Interestingly, Prx did not fit well into the modelled density possibly indicating that its conformation in solution could be different from that of crystal structure.

**Figure 6:**
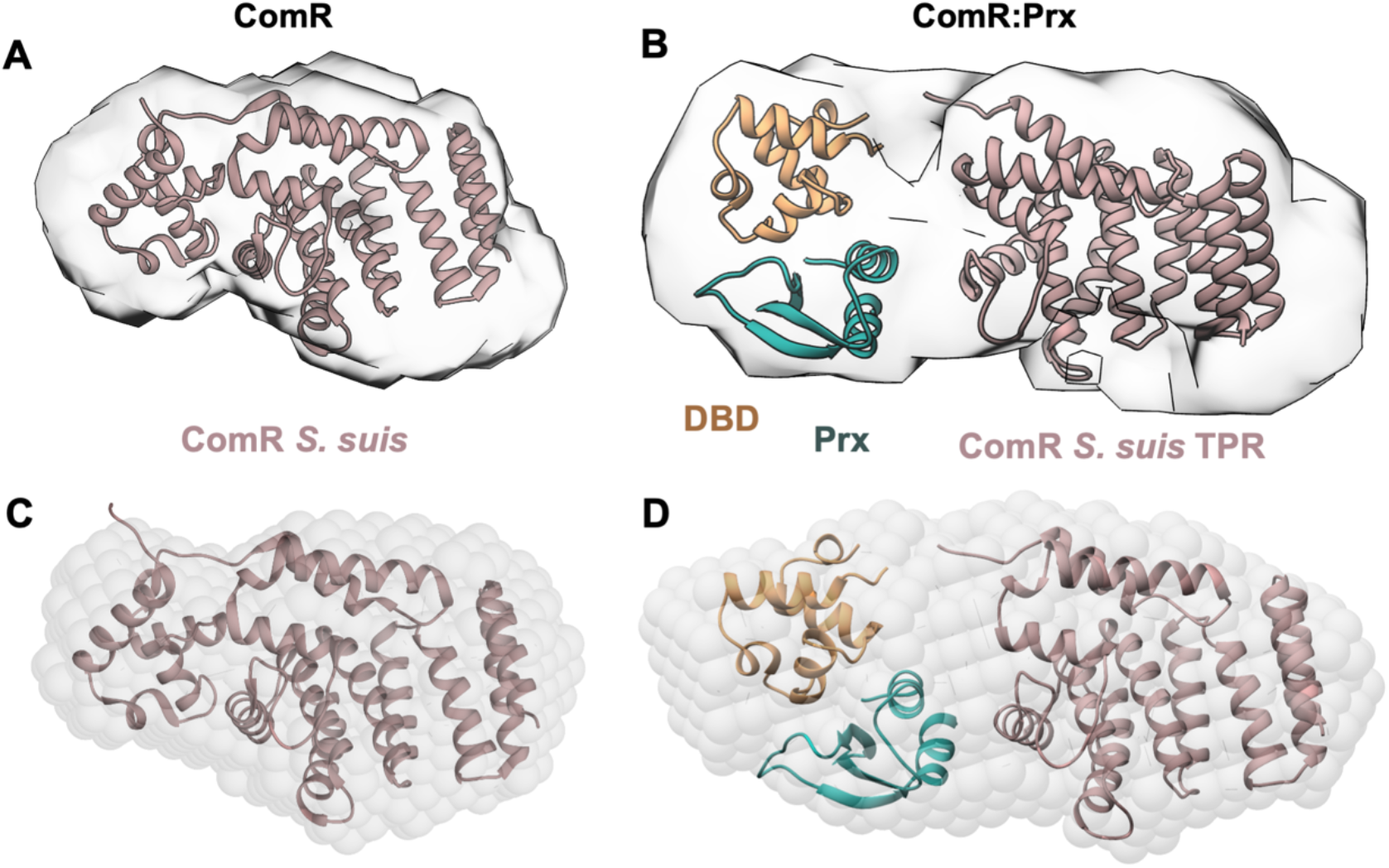
Low-resolution SAXS model of the full-length ComR:Prx complex. A) Full-length ComR (pink) (PDBid: 5FD4) modelled into a SAXS density envelope calculated by DENSS. B) ComR:Prx complex (orange:blue) modelled into a SAXS density envelope calculated by DENSS using the ComR *S. suis* TPR domain and a DBD:Prx crystal structure. C) As panel A but with a SAXS density envelope calculated using DAMMIN. D) As panel B but with a SAXS density envelope calculated using DAMMIN. All modeling was performed manually using Chimera.

### Prx inhibits ComR by preventing DNA binding independently of activation by XIP

Although the structural and biochemical data shows that Prx directly manipulates the conformation of ComR, the data does not address if Prx exerts an effect on the interaction of XIP with ComR. Additionally, since XIP induces a drastic conformational change to activate ComR is this conformational change in competition with the Prx induced conformational change? To address this question, we first performed electromobility shift assays (EMSAs) with ComR, XIP, Prx and FAM labelled *comS* promoter (Fig. 7). When ComR, XIP, and were Prx added together, Prx completely prevented ComR from binding DNA in a dose dependent manner (Fig. 7A left). This is similar to previous results, including the small observed shift of the ComR:XIP:DNA ternary complex in the lane 3 control (18). Next, we repeated this experiment but with either the pre-formation of the ComR:Prx complex or pre-formation of the ComR:XIP:DNA complex. When ComR and Prx were incubated together first followed by the addition of XIP and DNA, similar results were observed to when all components were added together (Fig. 7A middle). Specifically, we observe a dose dependent inhibition of the ability of ComR to bind DNA. Additionally, when Prx was added after the ComR:XIP:DNA complex was allowed to form Prx could still block the interaction of ComR with DNA (Fig. 7A right). Moreover, Prx appeared to be better at inhibiting the activity of ComR after activation by XIP. In this experiment, only 2 μM Prx was required to observe complete inhibition of activated ComR, or no DNA band shift, as compared to 6 μM Prx with apo-ComR. Identical experiments were completed with PrxD32A, which showed no inhibition in ComR:DNA complex formation. As residue D32 in Prx is absolutely essential for binding apo-ComR (Fig. 3A and Fig. 4), the observed inhibition of ComR by Prx is specific to direct interaction with the DBD.

**Figure 7.**
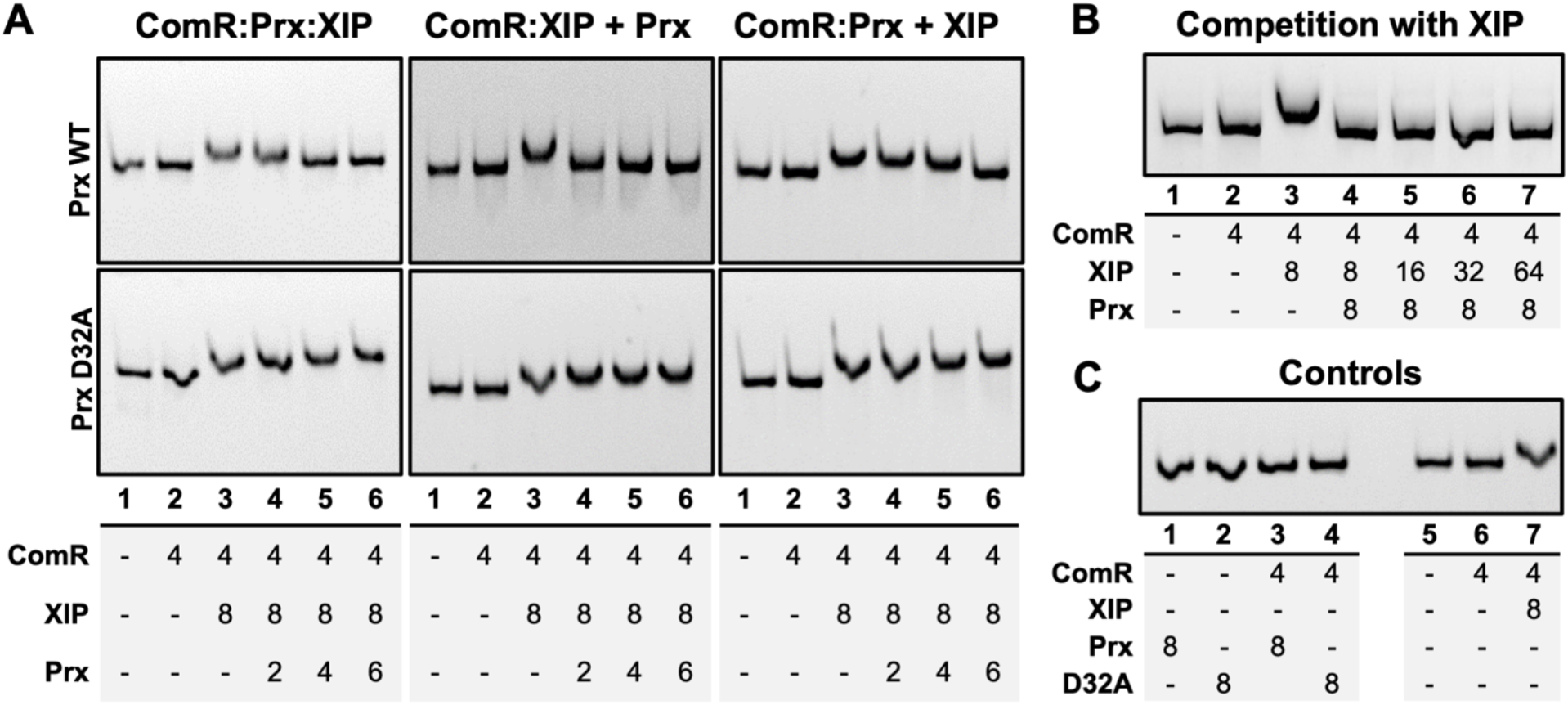
Electromobility shift assays of Prx with ComR:DNA complexes. A) (left) Experiment with ComR, Prx, and XIP incubated before the addition of the DNA probe. (middle) ComR was incubated with Prx followed by the addition of XIP and DNA. (right) ComR, XIP, and DNA were incubated to allow the ternary complex to form before the addition of Prx. Top panels are with wild-type Prx and bottom panels with the Prx D32A variant. B) ComR:Prx complexes challenged with an increasing molar excess of XIP C) Control experiments with each assay component with DNA, including the ComR:Prx complex and the ComR:XIP complex. A DNA shift can be seen only in the presence of ComR and XIP. Under each EMSA the individual experiments are numbered by gel-lane with the μM concentration of each component listed.

Given that Prx can completely inhibit ComR *in vitro,* we next asked if the interaction of XIP with ComR could out-compete Prx. As shown in Figure 7B, regardless of the amount of XIP added in the EMSA XIP could not overcome the ability of Prx to inhibit ComR. Specifically, even at an 8-fold and 16-fold molar excess of XIP relative to Prx and ComR respectively, ComR was unable to bind DNA (Fig. 7B). Additional controls with all EMSA components are shown in Figure 7C indicating that these observations are not due to nonspecific interactions with the DNA probe.

To further address if Prx is a competitive inhibitor of XIP activation, we performed binding assays using ComR and fluorescently labelled XIP with Prx. First, we measured the binding of dansyl-XIP to ComR. Dansyl-XIP has a low fluorescence quantum yield in solution, however upon binding ComR the fluorescence quantum yield significantly increases. This experiment yielded a K_d_ of 0.32 ± 0.04 μM (Fig. 8A and Fig. 8B) showing higher affinity to ComR in comparison to previously reported values determined by ITC for unlabelled XIP with ComR (18). This difference is not unexpected, given that the dansyl moiety could hydrophobically interact with residues in the XIP binding pocket and that ComR proteins can be sensitive to the exact length of the peptide (23). To verify that the interaction of dansyl-XIP with ComR was equivalent to that of unlabeled XIP, we tested if XIP could displace the fluorescent probe. As shown in Figure 8C, XIP competes with dansyl-XIP reducing the observed fluorescence in a dose dependent manner. We then repeated the same competition experiment but instead with Prx. Increasing concentrations of Prx added to ComR:dansyl-XIP showed minimal changes in fluorescence indicating that Prx is unable to displace the dansyl-XIP from ComR. This could either mean that Prx was unable to bind ComR when XIP is bound, or that Prx can bind the XIP activated conformation of ComR. As the former was highly unlikely given the EMSA data (Fig. 7), we assayed if Prx could form a complex with ComR:XIP. In order to test if Prx can interact with ComR:XIP we performed the same competition assay in Figure 8C, but instead of monitoring the fluorescence intensity, we have measured the steady-state fluorescence anisotropy of the ComR:dansyl-XIP complex. The anisotropy of a fluorescently labeled molecule in solution is dependent upon its rotational correlation time and can be used to monitor changes in its size and potential protein complex formation (37). Although Prx is a small 8 kDa protein, we were able to see an increase in fluorescence anisotropy with the addition of Prx to ComR:dansyl-XIP in a dose dependent hyperbolic fashion, showing that Prx binds the complex with a sub-micromolar affinity of 0.7 ± 0.3 μM (Fig. 8D). Together with the EMSA data, this suggests that the interaction of Prx with ComR is independent of that with XIP and ComR. Specifically, Prx can form a ternary complex with activated ComR:XIP and the Prx induced conformational change does not affect the ability of XIP to bind the ComR TPR domain.

**Figure 8.**
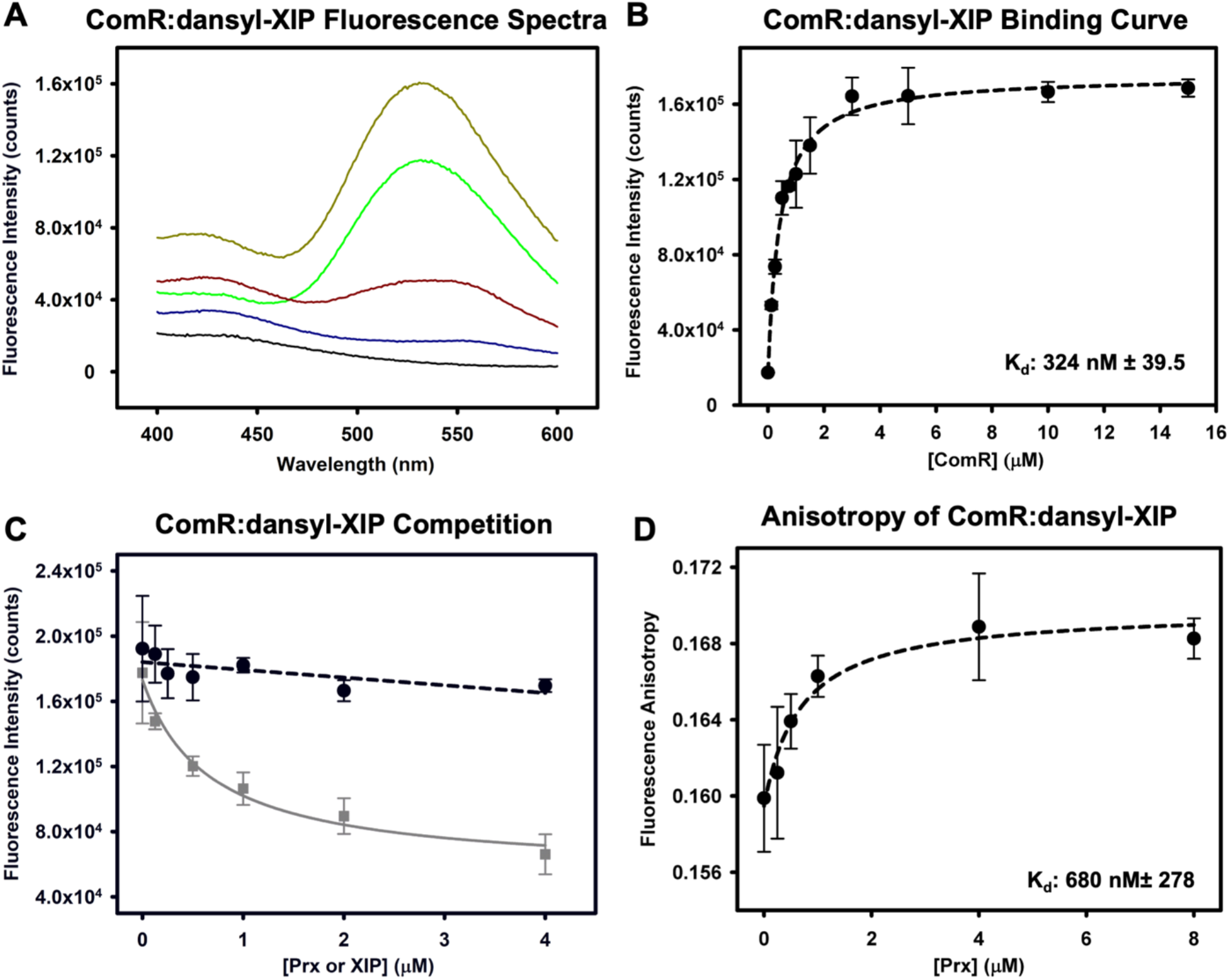
Prx binding to ComR does not affect the interaction with XIP. A) Example of observed fluorescence at the dansyl emission for increasing concentrations of ComR. Black is buffer background, blue is 0 μM ComR, red is 0.125 μM ComR, green is 0.75 μM ComR, and brown is 15 μM ComR. B) Binding curve of fluorescently labeled Dansyl-XIP with wild-type ComR *S. mutans.* A constant concentration of dansyl-XIP (0.2 μM) with increasing ComR was used as defined in the Methods section. C) Competition assays for dansyl-XIP bound to ComR with unlabeled XIP (grey) and Prx (black). Prx and XIP were added at the same molar ratios to the ComR:dansyl-XIP complex. Increasing XIP displaces dansyl-XIP and Prx has no effect. D) Fluorescence anisotropy measurements of Prx added to ComR:Dansyl-XIP demonstrating that Prx can bind the ComR:XIP complex. Error bars are representative of three separate experiments.

## DISCUSSION

Using a diverse set of biochemical and biophysical experimentation, our work has revealed a molecular mechanism of quorum sensing inhibition by bacteriophage in *Streptococcus pyogenes.* We show the specific structural manipulation of the quorum sensing receptor ComR by the prophage protein Prx and consequently how this small phage protein can inhibit natural transformation. Specifically, Prx induces a substantive conformational change in apo-ComR to bind directly to the normally protected DNA-interacting residues of the DBD (Fig. 3, Fig. 5, and Fig. 6). This in turn blocks association of ComR with its target promoter regions for *comS* and *sigX* (Fig. 7). Moreover, the interaction of Prx with ComR is independent of the interaction of XIP with ComR. Prx can bind the XIP activated conformation of ComR without displacing XIP (Fig. 8), and XIP cannot outcompete the ability of Prx to inhibit ComR (Fig. 7). This has the potential to be especially advantageous for inhibition of the quorum sensing circuit. Namely, the effective concentration of XIP in the cell would be reduced as it is rendered sequestered and ineffective while trapped in a ComR:XIP:Prx ternary complex. Overall, this is a primary example of the sophistication and ingenuity of bacteriophage to expertly manipulate their hosts.

Interestingly, we found that Prx is broad but specific to ComR as Prx can directly interact with Type-I, Type-II, and Type-III ComR proteins but appears to completely ignore the close structural homologue Rgg3 also found in *Streptococcus* (Fig. 1 and Fig. S1). The ability of Prx to bind different ComR variants was not surprising as the DBD:Prx crystal structure revealed that the phage protein targets residues conserved across all ComR proteins. However, given that ComR and Rgg3 belong to the same subfamily of structurally related RRNPP quorum sensing regulators, RRNPP selectivity was not immediately apparent. Both ComR and Rgg3 possess a DBD and TPR domain that recognizes a specific peptide pheromone (12). The DBD of ComR *S. mutans* and Rgg3 *S. pyogenes* are 24% identical with the same overall fold (RMSD backbone 1.4Å^2^). This includes residue conservation and structural overlap with the Prx interaction residues ComR(Q34) with Rgg3(Q32) and ComR(R37) with Rgg3(R35) (PDBid: 6W1A) (Fig. 3A). However, in Rgg3 T32 occupies the same structural position as R33 in the ComR DBD which Prx(D32) interacts with. Other variations like this exist which could disrupt the hydrogen bond network that appears to stabilize Prx bound to the ComR DBD. Furthermore, structural studies have shown that unlike apo-ComR, apo-Rgg3 is already a dimer before activation by pheromone and the DBDs are dynamic relative to the TPR domain (26). This difference could serve to further reduce the ability of Prx to bind Rgg3 through steric hindrance of already dimerized DBD.

To our knowledge, the molecular mechanism of inducing a large conformational change in a quorum sensing transcription factor by a phage protein is unique. Prx can bind to both apo-ComR and activated ComR conformations independently of the pheromone XIP to inhibit transcription of competence genes (Fig. 7 and Fig. 8). When targeting the apo-form of ComR, Prx specifically binds residue Q34 which helps unhinge the DBD from the TPR (Fig. 3) (23). Interestingly, our past results (18) and our SAXS data show that this complex does not dimerize with another ComR (Fig. 5 and Fig. 6). Furthermore, despite releasing the DBD, Prx does not appear to alter the conformation of the TPR as XIP is not displaced (Fig. 8). This is in contrast to the mechanism of XIP binding, in which XIP specifically manipulates the helices of the TPR to force release of the DBD (22, 23). Based on our structural data, the biochemical mechanism of quorum sensing inhibition by Prx is to both shield the DNA interacting residues and to uncouple the conformational change of the TPR from the DBD.

The ITC data (Fig. 2C and Fig. 2D) and the EMSA data (Fig. 7) suggest that Prx has a higher affinity for the activated form of ComR than apo-ComR. This is supported by the observed release of the DBD from the TPR upon XIP binding such that Prx would no longer be required to break the DBD:TPR interaction surface (23). Importantly, this allows Prx to both pre-empt signaling by its host and to stop the signaling cascade for natural competence once it has begun. Overall, since Prx binds to ComR with high affinity regardless of conformational state, this highlights an extreme biological importance of inhibiting natural competence, or at least ComR, by the phage. Specifically, this could be to serve in self-protection by the prophage from being damaged or replaced by recombination during natural transformation. However, deletion of *prx* from various *S. pyogenes* strains only decreases the expression of competence late genes without increasing the ability of *S. pyogenes* to actually transform (18). This still leaves several questions remaining as to what is also blocking natural competence and why the phage has specifically evolved to inhibit ComR.

While the structural details for the manipulation of ComR by Prx remains unique, an analogous mechanism of quorum sensing inhibition by a small bacteriophage protein was recently described in the Gram-negative bacteria *Pseudomonas aeruginosa* (20). The *Pseudomonas* phage DMS3 uses a small protein Aqs1 to inhibit the master quorum sensing regulator and transcription factor LasR. Overall, this prevents the expression of a number of genes ranging from motility to anti-phage defenses. Although both Prx and Aqs1 interact with the conserved DNA binding amino acids in their respective transcriptional regulators, the phage proteins share no structural homology. Prx in complex with the ComR DBD is a monomer with a mixed alpha-beta fold, while Aqs1 is a dimer of two helical hairpins (Fig. 9). Additionally, it should be pointed out that although ComR and LasR are both quorum sensing receptors they also share low structural homology (20, 23) (Fig. S7A). Despite these differences, both Prx and Aqs1 use electronegative surfaces to form ionic interactions with the electropositive surfaces of their respective transcriptional regulators (Fig. S7B). Furthermore, at the molecular level both phage proteins inhibit transcription by contacting residues in the DBD helix that participate in DNA binding (Fig. 9) (20). Taken together, this highlights a prominent example of convergent evolution between two distinct bacteriophages in the regulation of their hosts. Not only does this show that both Gram-positive and Gram-negative bacteriophages have both evolved mechanisms to inhibit quorum sensing in their hosts, but it further emphasizes the amazing versatility of phage proteins.

**Figure 9.**
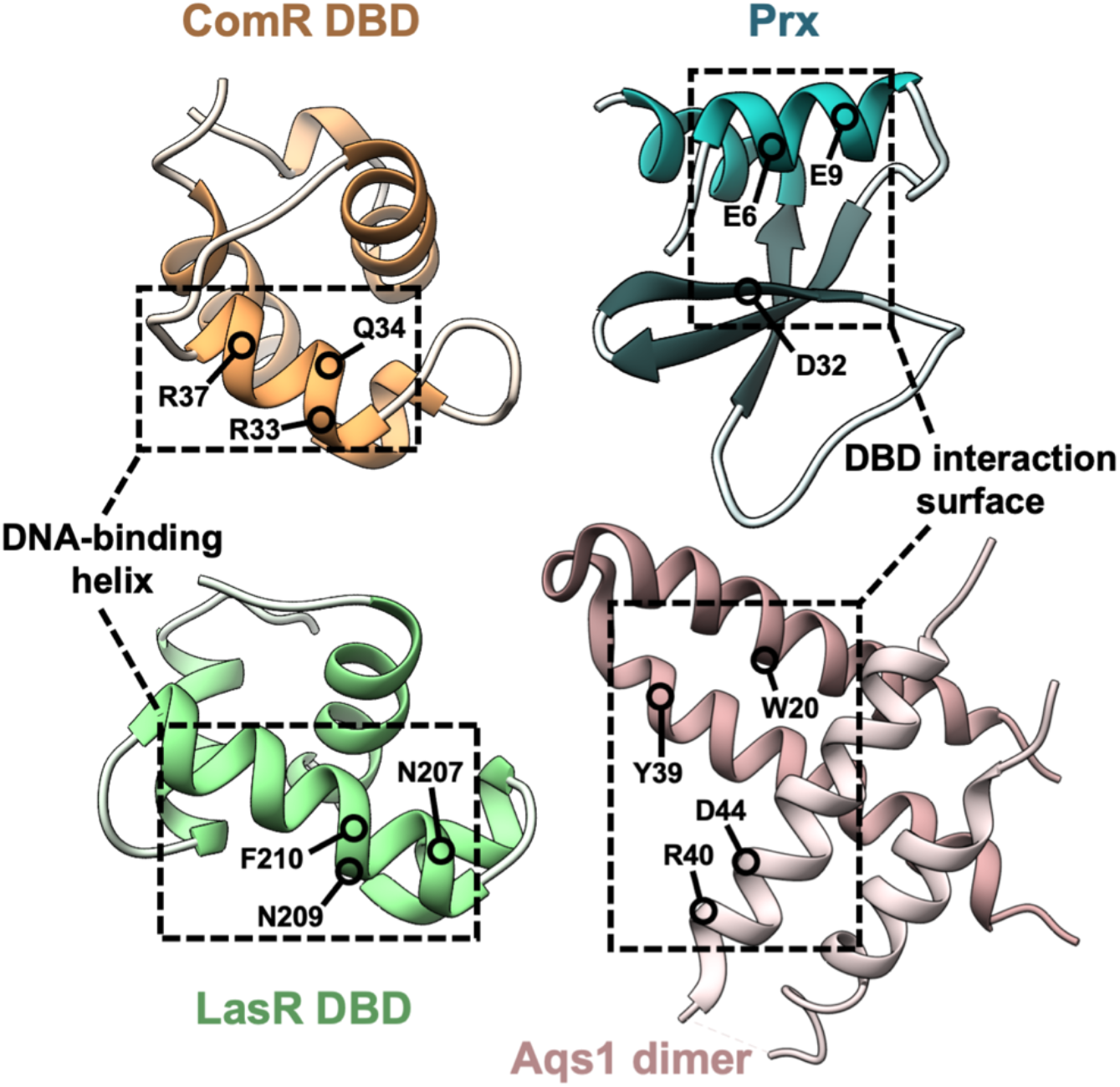
Comparison of the ComR:Prx complex and the LasR:Aqs1 complex. Left: The phage protein binding surfaces of the ComR DBD (orange) and the LasR DBD (PDBid: 6V7W) (green) are shown with the DNA-binding helix of each domain outlined. Residues that Prx and Aqs1 contact directly are indicated on each DBD respectively. Right: Prx (blue) and Aqs1 (PDBid: 6V7W) (pink) are displayed with the DBD binding surfaces outlined. Residues that make contacts with the DNA-binding helices are shown on each structure.

As Prx functions in a similar role to Aqs1, it raises the question if Prx has other similarities to the *Pseudomonas* phage protein. In addition to being able to bind LasR, Aqs1 uses a distinct surface to bind PilB (20). PilB is an ATPase required for type IV pilus assembly, which Aqs1 directly inhibits (38). As a type IV pilus is crucial for natural competence, does Prx play a similar dual role? Furthermore, the biological link between Prx and the phage toxins remains unknown. Given the precedent of small phage proteins to evolve multiple roles to economize on the small genome of the phage (20), it is reasonable to hypothesize that Prx has an additional role related to the transmission of phage toxins. Regardless of what additional biochemical activities remain to be discovered, it is clear that Prx uses a unique mechanism to block bacterial communication and protect the phage genome from natural transformation.

## EXPERIMENTAL PROCEDURES

### Protein expression and purification

The protein expression constructs used include Prx MGAS315 and ComR *S. mutans* (18), and ComR *S. suis* (23). Expression vectors containing ComR MGAS5005 and Rgg3 *S. pyogenes* were generously gifted by the Federle lab from the University of Illinois at Chicago, Chicago, IL USA. The expression construct for ComR *S. thermophilus,* was ordered, codon optimized in pET21a, from Genscript. For the ComR *S. mutans* DBD construct the coding sequence for ComR *S. mutans* was truncated with the addition of a stop codon after residue D66. by Q5 mutagenesis (New England Biolabs). For the TRR construct, primers TPR_NdeI_F and TPR_BamHI_R were used to amplify the TPR domain (residues S74 to T304) of ComR *S. mutans* and the product was cloned into the NdeI and BamHI sites of an empty pET15b vector. Each Prx protein variant was generated through Q5 mutagenesis (New England Biolabs). The described primers were ordered from Integrated DNA technologies (IDT) and are listed in table S1. All expression vectors were transformed into *E. coli* BL21(DE3) Gold cells for protein expression.

For the purification of each ComR ortholog, Prx and variants, ComR DBD, and the ComR TPR domain, proteins were expressed and purified as previously described (18). To summarize, cells were grown in LB + 100 μg/mL ampicillin at 37 °C to an optical density at 600 nm of 0.6 to 0.8, at which point the temperature was reduced to 20 °C and isopropyl β-D-1-thiogalactopyranoside (IPTG) was added to a concentration of 1 mM to induce protein expression. The cells were collected the following morning by centrifugation, and the cell pellets were re-suspended in wash/lysis buffer (50 mM Tris-HCl pH 7.5, 500 mM NaCl, 25 mM Imidazole), and stored at −80 °C. Before lysis, cell pellets were thawed and 1 mM PMSF, 10 mM MgCl2 and a small amount of DNaseI were added. The cells were lysed with an EmulsiFlex-C3 (Avestin), and the soluble protein was separated by centrifugation at 16000 rpm. The soluble lysate was then purified over a nickel-NTA gravity column, washed with 250 mL of wash/lysis buffer, and eluted with elution buffer (50 mM Tris-HCl pH 7.5, 500 mM NaCl, 500 mM imidazole). Lysate and/or protein samples were kept on ice or at 4 °C between each purification step.

For each ComR protein, the His-tag was removed by dialysis overnight at 4 °C in gel filtration buffer (20 mM Tris-HCl pH 7.5, 100 mM NaCl, 1 mM β-ME) at an approximate molar ratio of 100:1 ComR:thrombin (for ComR *S. mutans*, *S. suis*, and MGAS5005) or ComR:HRV 3C Protease (for ComR *S. thermophilus*). The digested protein was re-run over the nickel-NTA column and the flow-through was collected. SUMO-Rgg3 was also purified as described and incubated with SUMO protease overnight at 4 °C. An approximate molar ratio of 80:1 protein:enzyme was used in a low-salt digestion buffer (50 mM Tris pH 7.5, 100 mM NaCl, 2 mM β -ME and 10 % glycerol).

Following affinity chromatography and/or affinity tag removal, each protein was further purified, by size exclusion chromatography (SEC), allowing for complete buffer exchange into gel filtration buffer. Proteins were concentrated and run over a HiLoad 16/600 superdex 75 gel filtration column (GE Healthcare) using an AKTA pure (GE Healthcare).

### Size exclusion chromatography binding assays

For each binding assay, proteins were incubated on ice for 15 min, at a 1:1.5 molar ratio of ComR:Prx, with the exception of the DBD, where a higher molar ratio of DBD was used. Protein complexes were run over a HiLoad 16/600 superdex 75 column (GE Healthcare) or a Superdex75 increase 10/300 column (GE Healthcare) using an AKTA pure (GE Healthcare). All assays were performed with gel-filtration buffer (20 mM Tris-HCl pH 7.5, 100 mM NaCl, 1 mM β-ME).

### Pull-down assays

Ni-NTA resin (GoldBio) was prepared by washing (addition of water/buffer, incubating/rotating for 5 min at 4 °C, centrifugation at 1500 × g for 5 min, removal of supernatant) twice with water and then with buffer (20 mM Tris-HCl pH 7.5, 100 mM NaCl, 1 mM β -ME). 20 μL of the prepared resin was added into individual centrifuge tubes, and a total of 0.36 mg of Prx (with a His-tag) was added to each. Following incubation for 30 min at 4 °C, and two wash steps with buffer, 40 μL at 1 mg/mL (0.4 mg total) of each ComR ortholog (without a His-tag) was added to individual Prx saturated resin samples. Protein/resin mixtures were mixed for 1 h at 4 °C, then washed twice with buffer. Next 15 μL of 4× Laemmli sample buffer was loaded directly into each sample which was then boiled at 93°C for 5 min, and centrifuged at 21000 × g for 3 min. 5 μL of each sample was then run on a 15 % SDS-PAGE gel. The No-Prx controls (ComR background) were performed alongside and identically to the pull-downs, with buffer added rather than Prx. ComR only controls, of similar amounts of the ComR added to the pull-down experiment, were also run on an SDS-PAGE gel to ensure sample purity and consistent protein concentration measurements.

### Isothermal titration calorimetry

Each protein was dialyzed overnight at 4 °C in the same buffer (gel filtration buffer). ITC was performed using a MicroCal iTC200 (GE Healthcare). 300 μM Prx was injected into 20 μM ComR *S. mutans* or DBD at a constant temperature of 25 °C. Controls of Prx injected into buffer, as well as buffer injected into both ComR and the DBD were performed. The same experiment was repeated with 300 μM PrxD32A injected into 20 μM ComR. Data analysis was performed with the Origin software (Malvern) and the results are based on a one site model.

### Protein Crystallization

For protein crystals of the DBD:Prx complex, proteins were incubated at 1.5:1 molar ratio of DBD:Prx and run over the HiLoad 16/600 superdex 75 size exclusion column. The fractions containing the complex were collected, run on an SDS-PAGE gel to assess purity, and concentrated. Protein crystallization was screened for in commercially available screens (Qiagen), using a Crystal Gryphon robot (Art Robbins Instruments). Crystals were grown in 1:1 gel filtration buffer with 15 mg/mL and 0.2 M Potassium acetate, 20 % PEG 3500. Coincidentally, protein crystals of the DBD:Prx complex also formed as a result of *in situ* proteolytic digestion of the ComR:Prx complex using the Proti-Ace kit (Hampton). Crystals formed after 10 mg/mL ComR:Prx complex was digested with 50 μg/mL α-chymotrypsin at 4 °C for 1 h. The digested protein was screened for crystallization in a variety of commercially available screens and crystals formed in 1:1 gel filtration buffer and 0.1 M HEPES pH 7.0 and 18 % PEG 12000. Each crystallization condition was screened for by sitting vapor-drop diffusion at 4 °C.

### Data collection & refinement

Datasets were collected at the Advanced Light Source (ALS) Beamline 8.3.1, as well as the Canadian Light Source beamline CMCF-ID (081D). Protein crystals were first cryo-protected in each of crystallization screen conditions, along with the addition of the appropriate PEG (PEG 3350 or 12000) to a concentration of 35 %, and then flash frozen in liquid Nitrogen. Both of the datasets were processed using XDS (39) and CCP4 (40). The phases were solved by molecular replacement with an existing Prx structure (PDBid: 6CKA (18)) and the DBD of a structure of ComR *S. suis* (PDBid: 5FD4 (23)) using Phenix (41). The structures were built in Coot (42) and further refined using Phenix (41), Refmac5 (43), and TLS refinement (44).

### Circular dichroism

Concentrated proteins of Prx MGAS315 (PrxWT), PrxE6A, PrxE9A, PrxD12A, PrxF31A, PrxD32A were dialyzed in 1.5 L of filtered CD buffer (20 mM potassium phosphate pH 7.5, 100 mM NaF) at 4 °C overnight. The following morning each protein was diluted to an appropriate stock concentration using the CD dialysis buffer. The CD was performed using a Jasco J-810. Each protein was at a concentration of 0.2 mg/mL, and loaded into a 0.05 cm cell. Three accumulations of each sample, from 260 nm to 190 nm, were collected and averaged.

### Small angle X-ray scattering

SEC-SAXS data for ComR, ComR:Prx, and Prx, were collected at 9mg/mL at the B21 beamile, Diamond light source (UK). Using a Shodex KW402.5-4F column equilibrated in 20 mM Tris-HCl pH 7.5, 100 mM NaCl, 1 mM β-Me at the flow rate of 0.160 mL/min. Each frame was exposed to the X-rays for 3 seconds, as described previously (45).Data processing and analysis was first performed using the ATSAS (46) software package, including CHROMIXS (29) for peak selection and buffer subtraction, and PRIMUS (30) for further data processing, including Guinier analysis and Kratky analysis. GNOM (33) was used for the calculation of a P(r) distribution and the D_max_ for both ComR and ComR:Prx. The buffer-subtracted datasets were processed identically in ScÅtter (www.bioisis.net) for the estimation of the Porod exponent (P_E_), providing a semi-quantifiable unit for protein flexibility, as well as the Porod volume (V_P_). Low-resolution electron density maps for ComR and ComR:Prx were estimated using DENSS (34). The processed SAXS data for each protein sample 20 density maps were calculated and averaged, then refined back to the original data set with DENSS (34). We also calculated 12 low-resolution models using the program DAMMIN (35). The individual DAMMIN models were rotated, averaged and refined to obtain a filtered model using the program DAMAVER (47), as previously (48). The protein structures, including ComR *S. suis* (PDBid: 5FD4) and DBD:Prx, were aligned to the density maps in Chimera (36).

### Electromobility shift assays

Fluorescently labeled DNA probes were amplified using the UA159 PcomS F, and FAM UA159 PcomS R primers, used previously (18). Electromobility shift assays were also performed as previously described (18), with 4 μM ComR, 8 μM XIP and increasing Prx, followed by the addition of 100 ng DNA probe. Each protein was added together (Prx, XIP, then ComR) and incubated for 30 min, flowed by the addition and incubation with DNA for 15 min. Additionally, the following experiments with the same protein concentrations were performed: pre-formed Prx:ComR complex, and pre-formed ComR:XIP:DNA. For the pre-formed Prx:ComR complex, Prx and ComR were first incubated for 30 min, followed by the addition of XIP and DNA for 15 min. For the next experiment ComR:XIP:DNA were incubated together for 30 min, followed by the addition of Prx, incubated for 15 min. Each of these experiments were also performed with a PrxD32A control. For the XIP competition experiment, ComR (4 μM), Prx (8 μM) and DNA (100 ng) were incubated together for 30 min, followed by the addition of increasing amounts of XIP. For each experiment, samples were run on a 5 % native PAGE gel at 100 V for 50 min at 4 °C, and imaged using a Fluorochem Q imager (Protein Simple).

### Fluorescence & Anisotropy binding assays

Both ComR *S. mutans*, and Prx were equilibrated in a final buffer of 20 mM HEPES pH 7.5, 100 mM NaCl, 1 mM β-ME, by size exclusion chromatography. The fluorescently labelled *S. mutans* XIP (dansyl-GLDWWSL) was stored at −80 °C and dissolved in the final buffer immediately prior to use. Pure, unlabelled *S. mutans* XIP (GLDWWSL) was also and dissolved in final buffer before use. Both peptides were synthesized by Abclonal. For the ComR:dansyl-XIP binding experiment, 0.2 μM dansyl-XIP was incubated with increasing amounts of ComR for 1 h, at room temperature, and in the dark. The following competition and fluorescence anisotropy experiments were performed with 0.2 μM dansyl-XIP, 0.5 μM ComR, and increasing amounts of either unlabeled XIP or Prx. Proteins and XIP were added together and incubated for 1 h at room temperature, in the dark. All fluorescence emission spectra were collected using a Jobin-Yvon Fluorolog-3 fluorescence spectrophotometer (Horiba) using 335 nm excitation light and scanned from 400 to 600 nm. For measuring steady-state emission spectra, the excitation and emission slit widths were set at 5 nm bandpass, while for anisotropy measurements they were set to 10 nm bandpass.

## ABBREVIATIONS

β-ME: 2-Mercaptoethanol
CD: Circular dichroism
DBD: DNA binding domain
EMSA: Electromobility shift assay
IPTG: Isopropyl β-d-1-thiogalactopyranoside
ITC: Isothermal titration calorimetry
PMSF: phenylmethylsulfonyl fluoride
SAXS: Small angle X-ray scattering
SEC: Size exclusion chromatography
TPR: tetratricopeptide repeat
XIP: SigX inducing peptide

## DATA AVAILABILITY

The X-ray structures and diffraction data reported in this paper have been deposited in the Protein Data Bank under the accession codes 7N10 and 7N1N.

## SUPPORTING INFORMATION

This article contains supporting information.

## ACKNOWLEDGEMENTS

We would like to thank the laboratory of Sean A. McKenna at the University of Manitoba for access to his Fluorochem Q imager for EMSA assays and the laboratory of Joe O’Neil at the University of Manitoba for access to his Jasco J-810. We thank Brian L. Mark at the University of Manitoba and Beamline 8.3.1 at the Advanced Light Source for assistance with X-ray data collection. We also thank beamline CMCF-ID at the Canadian Light Source, which is supported by the Canada Foundation for Innovation (CFI), the Natural Sciences and Engineering Research Council (NSERC), the National Research Council (NRC), the Canadian Institutes of Health Research (CIHR), the Government of Saskatchewan, and the University of Saskatchewan. Finally we thank the Diamond Light Source (UK) and the B21 Beamline scientists (SM22113) for SAXS data collection.

## FUNDING AND ADDITIONAL INFORMATION

This work was supported by the Natural Sciences and Engineering Research Council of Canada (NSERC) grants RGPIN-2018-04968 to G.P., a Manitoba Medical Services Foundation Operating Grant (MMSF) 8-2020-10 to G.P., and a Canadian Foundation for Innovation award (CFI) 37841 to G.P. TRP acknowledges Canada Research Chair program.

## CONFLICT OF INTEREST

The authors declare that they have no conflicts of interest with the contents of this article.

